# A convolutional neural network for common coordinate registration of high-resolution histology images

**DOI:** 10.1101/2020.09.18.303875

**Authors:** Aidan C. Daly, Krzysztof J. Geras, Richard A. Bonneau

## Abstract

Registration of histology images from multiple sources is a pressing problem in large-scale studies of spatial -omics data. Researchers often perform “common coordinate registration,” akin to segmentation, in which samples are partitioned based on tissue type to allow for quantitative comparison of similar regions across samples. Accuracy in such registration requires both high image resolution and global awareness, which mark a difficult balancing act for contemporary deep learning architectures. We present a novel convolutional neural network (CNN) architecture that combines (1) a local classification CNN that extracts features from image patches sampled sparsely across the tissue surface, and (2) a global segmentation CNN that operates on these extracted features. This hybrid network can be trained in an end-to-end manner, and we demonstrate its relative merits over competing approaches on a reference histology dataset as well as two published spatial transcriptomics datasets. We believe that this paradigm will greatly enhance our ability to process spatial -omics data, and has general purpose applications for the processing of high-resolution histology images on commercially available GPUs.

## 1 Introduction

In the study of tissue pathology, spatial context matters. Tissues are composed of a variety of distinct cell types, often arranged in complex spatial patterns and interacting with neighbors via an intricate signalling network. As a result, measurements of cellular quantities of interest, such as mRNA or protein, may have very different interpretations depending on the precise location from which they were drawn. For this reason, bulk or single-cell methods for measuring the transcriptome or proteome, which can attain high yield of RNA/protein targets but require dissociation of the tissue, may not be ideal due to the substantial amount of contextual information they discard [Wang et al., 2009, Hwang et al., 2018]. While computational approaches have been developed to map single-cell measurements back to coordinates in the original tissue [Edsgard et al., 2018, Stuart et al., 2019], recently developed high-throughput techniques for obtaining spatially resolved measurements of the transcriptome of *intact* tissue sections — either through highly multiplexed fluorescent *in situ* hybridization (FISH) [Sansone, 2019, Xia et al., 2019] or solid-phase mRNA capture [Ståhl et al., 2016, Rodriques et al., 2019] — provide an attractive alternative due to their enhanced ability to observe pathological mechanisms at sub-cellular scales within their native environment. As technical advances continue to increase both their throughput and resolution [Vickovic et al., 2019, Stickels et al., 2020, Asp et al., 2020], these so-called spatial “-omics” methods promise an unprecedented view of complex tissue function and dysfunction.

While spatial -omics measurements of single tissue sections can be informative for applications such as patient-centered medicine, detecting trends at the level of organ, individual, or even population requires integrated analysis of data from multiple sources. Due to biological and technical variation, however, this cannot be accomplished by a simple overlay of results, and requires an initial *registration* step: a transformation that maps coordinates in one system to corresponding coordinates in another. When registering sequential images of a single tissue, one can rely on cellular landmarks, such as DAPI-stained nuclei, to define homography transforms that either minimize distance or maximize mutual information between images. When registering image data from multiple sources, we cannot expect to define a direct homography between samples, and instead define registration in terms of correspondence of higher-level features. One such approach involves segmenting tissue into distinct anatomical annotation regions (AARs) corresponding to conserved tissue types, effectively registering tissue to a common coordinate system and allowing for the comparison of like regions across samples [Rood et al., 2019, Wang et al., 2020]. This approach was employed by Maniatis et al. [2019] in their spatial transcriptomics (ST) study of amyotrophic lateral sclerosis (ALS) in mice and humans. The authors employed the ST methodology of Ståhl et al. [2016], a solid-phase capture approach which yields unbiased measurements of the transcriptome at discrete capture areas spaced regularly across the tissue. These capture areas, or “spots,” are 100 μm in diameter, spaced 200 μm apart, and can each capture thousands of distinct mRNA reads. Subsequent iterations of this technology, such as 10X Genomics’ Visium platform, have attained greater spatial resolution (55 μm-wide spots, 100 μm center-center distance) and capture efficiency (10s of thousands of reads depending on tissue type) but operate on the same principle. By manually assigning each spot an AAR label based on visual inspection, the authors were able to effectively align data from multiple individuals and model gene expression in a hierarchical fashion conditioned on sample covariates (Fig. 1a). With an appropriately designed coordinate system, a similar registration strategy can be applied to other tissues, even those without stereotyped anatomy (pathologists routinely segment tumor biopsies by cancer state), to lend statistical power to a wide range of spatial -omics studies.

**Figure 1:**
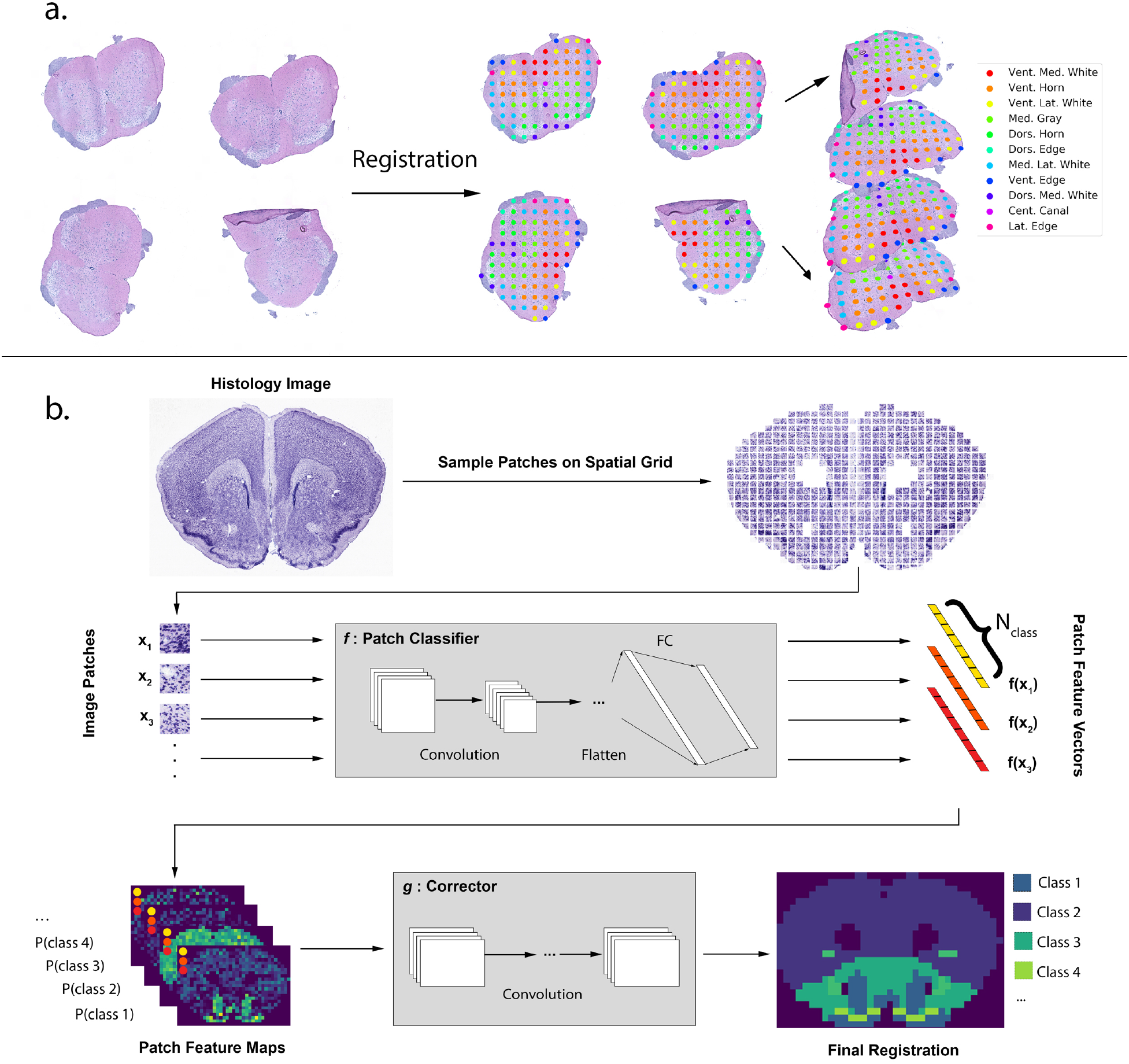
Common coordinate registration and the automation thereof by GridNet. a) An example of common coordinate registration using mouse spinal cord tissue and 11 tissue classes. b) Schematic of the proposed CNN architecture operating on a cross-section of mouse brain tissue. An image classification CNN *f* is employed to extract feature vectors from patches sampled from high-resolution histology images. These feature vectors serve as input to a second CNN *g* which is trained to perform semantic segmentation.

As the volume of data generated by spatial -omics studies increases, reliance on manual annotation quickly becomes infeasible. Consequently, deep-learning based approaches to image registration become attractive alternatives for their potential to speed throughput and enable large-scale experimental designs. Histological image data, however, present several unique challenges that merit consideration may hamper the application of existing deep learning approaches. Since tissue type exists on a continuum, AAR-style labels are difficult to assign with pixel-precision, and are more naturally applied to discrete areas (e.g, ST spots). This complicates the application of most segmentation architectures, even those like U-Net [Ronneberger et al., 2015] which have been designed for biological applications, which output pixel-level segmentations. While the sub-cellular resolution attained by FISH-based spatial -omics lends itself more naturally to pixel-level registration of individual cells, these experiments can scale poorly to large tissues (greater than 1cm^2^) due to lengthy image acquisition steps. Furthermore, the size of the tissue images being considered for either solid-phase capture or FISH-based methods — potentially tens of millions of pixels per image — limits our ability to process whole images using deep segmentation architectures on commercially available GPUs. While reduction of network complexity or down-sampling of images can help ease memory burden, such changes may result in the loss of ability to detect key sub-cellular features. An alternative approach that preserves these features relies instead on *sub*-sampling the input image at regions of interest (ROI), such as ST spot locations, and using a convolutional neural network (CNN) to independently classify each one. While this local approach solves the problem of input size, and was employed by Maniatis et al. [2019] in their development of the annotation tool Span,^1^, it is highly susceptible to local in the absence of information on spatial context of patches. This can be observed in the results of Tan et al. [2020], who apply a similar CNN architecture to the task of identifying cancer state in dissociated image patches from an ST study of prostate cancer biopsies. While some of these errors could be corrected by the application of denoising strategies for image segmentation such as conditional random fields [Lafferty et al., 2001], a more powerful approach would be able to learn characteristics of tissue organization, and correct local predictions with a birds-eye view.

Here, we present “GridNet”, a novel deep learning architecture for the registration of histology images to a common coordinate system. This network achieves an ability to model complex data by combining two CNNs: a classifier that is applied to full-resolution image patches extracted at sparsely sampled locations in the original histology image, and a segmentation network that operates on feature vectors extracted from each patch (Fig. 1b). This network can be trained end-to-end, with the classification CNN learning cellular and sub-cellular features key to identifying tissue region and state, and the segmentation CNN learning global tissue patterns to correct local errors. We explore several strategies for training GridNet, demonstrating approaches that address the differing demands of the component networks and are capable of being executed on a single GPU. We apply GridNet, as well as competing architectures for common coordinate registration, to two datasets with available gold-standard manual labels: a mouse brain section reference histology dataset obtained from the Allen Brain Atlas (ABA) [Wang et al., 2020], and the mouse spinal cord ST dataset detailed in Maniatis et al. [2019]. Using these data, we show that pure-segmentation and pure-classification approaches suffer, respectively, from limitations on model complexity and persistent local errors, while GridNet is able to leverage elements of both paradigms to attain consistently high registration accuracy. Finally, we apply GridNet to a dataset of human dorsolateral prefrontal cortex (DLPFC) tissue processed with 10x Genomics’ newly released Visium spatial transcriptomics platform [Maynard et al., 2020], which achieves high resolution by hexagonal packing of ST spots, in order to demonstrate the generality of our approach to new spatial systems. We believe our hybrid approach, for which we provide a publicly available Python implementation, presents a useful paradigm for general tasks in large-scale histology image processing, due to its demonstrated ability to blend of information from full-resolution local features and global tissue patterns.

## 2 Materials and Methods

### 2.1 Data Specification

In this section, we discuss the manner by which image data were collected for this study, and how the points on the “registration grid” from which we extract full-resolution image patches were chosen. For each tissue, image patches are sampled at the points in this grid and arranged into a five-dimensional array — height of registration grid, width of registration grid, height of patch, width of patch, number of image channels — that serves as an input to our tissue registration neural networks.

#### 2.1.1 Allen Brain Atlas (ABA) reference histology

The ABA dataset, summarized in Fig. 2, was constructed from 235 Nissl-stained sequential coronal sections of the mouse brain, which are publicly available at https://connectivity.brain-map.org/static/referencedata. For each image, we sampled patches from a 2-dimensional cartesian grid across the image, with a center-to-center distance of 192 pixels. This corresponds to approximately 200 μm, which simulates the center-center distance in the 10x genomics standard ST array used by Maniatis et al. [2019]. This sampling strategy required a 49 × 33-sized grid to span the tissue area of the largest section, and as such these dimensions were chosen as the fixed size of our simulated ST array. Using each position in the array as a centroid, we sampled patches of either 128 × 128 pixels or 256 × 256 pixels, depending on the input size of the architecture. For all patches, each image channel was separately normalized according to *x_c_* = (*x_c_* – *μ_c_*)/*σ_c_*, where *μ_RGB_* = (0.485, 0.456, 0.406), *σ_RGB_* = (0.229, 0.224, 0.225).

**Figure 2:**
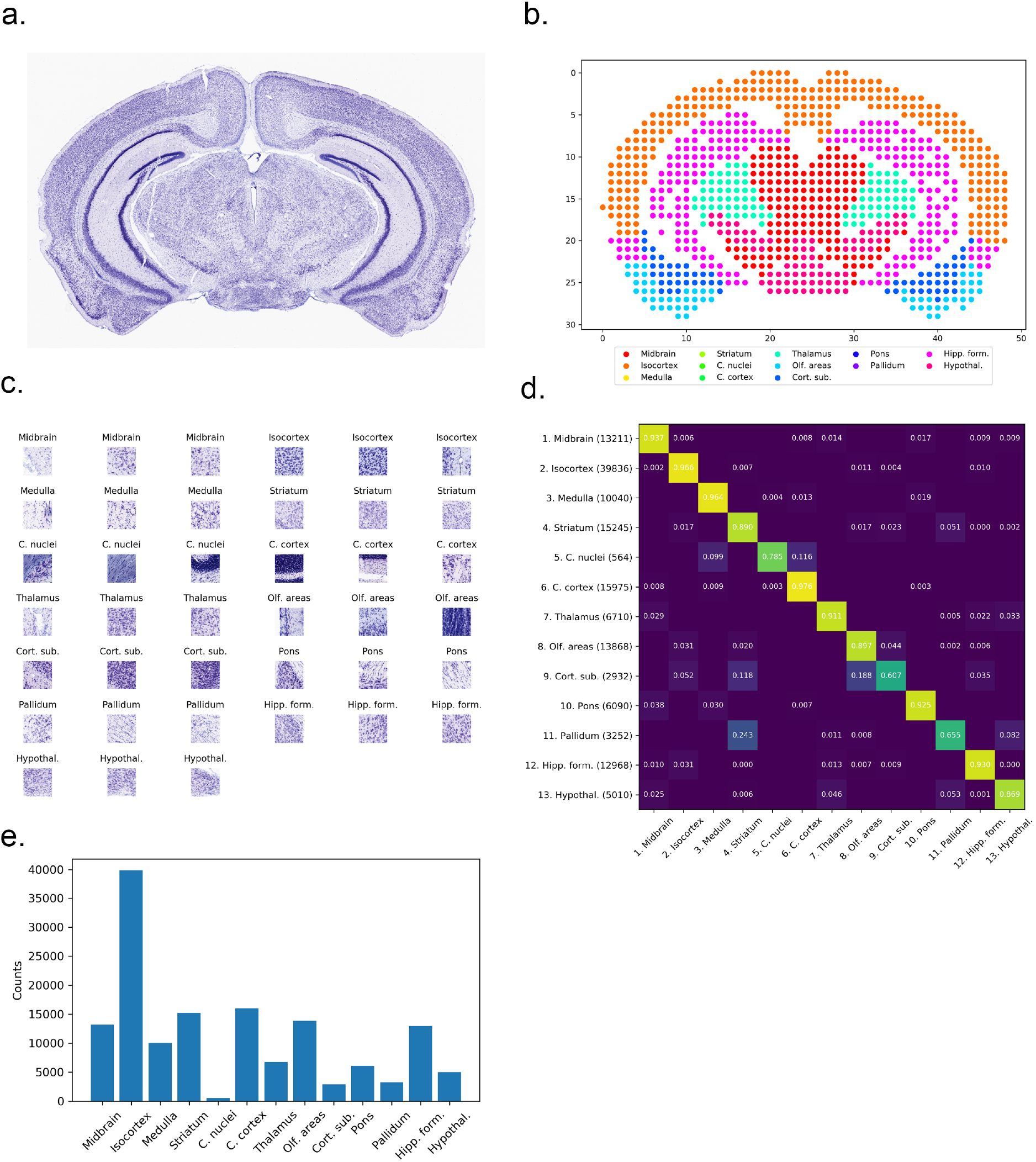
ABA reference histology dataset. (a) Representative histology image depicting a Nissl staining of a mouse brain coronal section. (b) Scatterplot displaying locations of corresponding ST spot centroids colored according to AAR annotation. (c) Three representative patches from each ontology region. (d) Adjacency matrix displaying the frequency with which patches of class [row] neighbor patches of class [column]. The number of instances of each class in the dataset is indicated in parentheses next to each row label in (d), as well as being displayed graphically in (e).

For each patch, an annotation was obtained by using the ABA’ Image Synchronization API ^2^ to map the pixel coordinates of the patch centroid either to coordinates within a structural atlas of the mouse brain (http://atlas.brain-map.org/atlas?atlas=2) or to the slide background. Each position within the structural atlas is associated with a hierarchy of ontology terms, describing the location with increasing degrees of specificity. We chose the level-5 ontology term as label for each foreground patch, yielding 13 unique classes across the dataset. The complete dataset consisted of 149,943 foreground patches with 13 unique annotations across the 235 arrays. The number of patches belonging to each class is detailed in Fig. 2(d). For experiments exploring optimal training strategies for GridNet, the patch size was set to 128 pixels, with 80% of the arrays used for training and 20% used for validation. For experiments comparing GridNet against competing methodologies for registration, the patch size was set to 256 pixels, with 70% of arrays were used for training, 10% for validation, and 20% for testing.

#### 2.1.2 Maniatis mouse spinal cord ST

The Maniatis dataset, summarized in Fig. 3, was constructed from 416 whole-slide images of haematoxylin and eosin (HE)-stained cross-sections of mouse spinal cord. Each of these imaged tissues had been processed using the Spatial Transcriptomics workflow described in Salmen et al. [2018], which employs 100 μm-diameter mRNA capture probes arranged in a cartesian grid at 200 μm center-to-center distance. The dimensions of all ST arrays are 35 rows by 33 columns.

**Figure 3:**
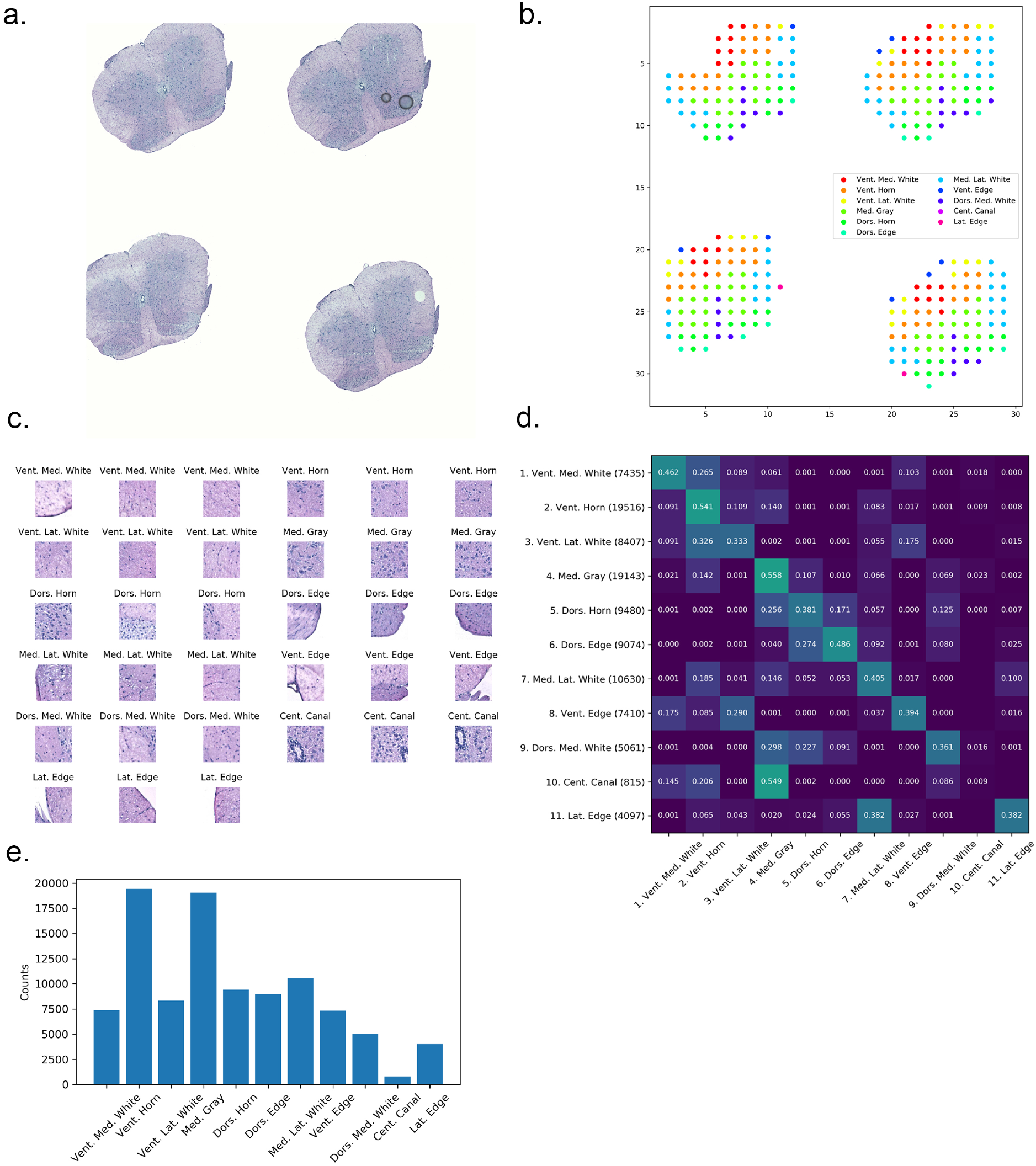
Maniatis mouse spinal cord ST dataset. (a) Representative histology image depicting hematoxylin and eosin (HE) staining of mouse spinal cord cross-section. (b) Scatterplot displaying locations of corresponding ST spot centroids colored according to AAR annotation. (c) Three representative image patches from each AAR, each displaying a (150*μm*)^2^ region. (d) Adjacency matrix displaying the frequency with which patches of class [row] neighbor patches of class [column]. The number of instances of each class in the dataset is indicated in parentheses next to each row label in (d), as well as being displayed graphically in (e).

Annotation files for each tissue section specifying the spatial location of each “foreground” ST spot — spots which overlapped tissue area and attained a minimum number of mRNA reads — along with the corresponding manually-assigned tissue labels (AARs) were obtained from Maniatis et al. [2019]. Using these foreground ST spot locations as centroids, patches were sampled to cover a 256 × 256 pixel region, which corresponded to a physical area of approximately 184 μm × 184 μm in each tissue. Due to variation in image resolution as a result of the data being collected from several distinct workflows, the width of the physical window size varied with a standard deviation of 34.5 μm across the dataset. For all patches, each image channel was separately normalized according to *x_c_* = (*x_c_* – *μ_c_*)/*σ_c_*, where *μ_RGB_* = (0.485, 0.456, 0.406), *σ_RGB_* = (0.229, 0.224, 0.225). The complete dataset consisted of 101,068 foreground patches with 11 unique classes across the 416 arrays. The number of patches belonging to each class is detailed in Fig. 3(d). For all experiments, 70% of the arrays were used for training, 10% for validation, and 20% for testing.

#### 2.1.3 Maynard human DLPFC Visium

The Maynard dataset, summarized in Fig. 4, was constructed from of 12 whole-slide images of HE-stained cross sections of human dorsolateral prefrontal cortex collected across three neurotypical patients. Each of these imaged tissues had been processed using 10x Genomics’ Visium Spatial Transcriptomics workflow, which employs 50 μm-diameter mRNA capture probes arranged in a hexagonal grid at 100 μm center-to-center distance. The dimensions of all Visium array are 78 rows by 64 columns.

**Figure 4:**
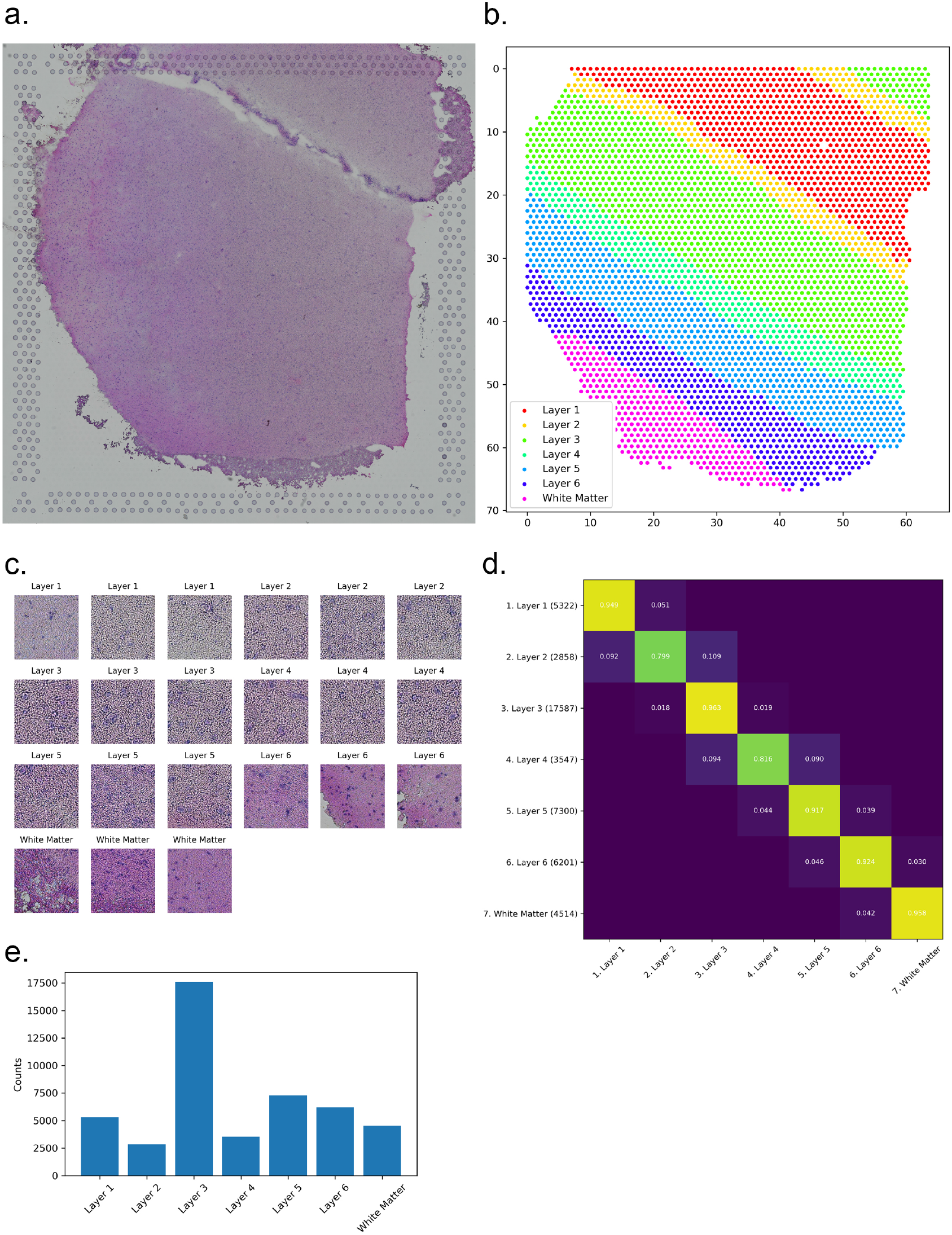
Maynard human dorsolateral prefrontal cortex (DLPFC) Visium ST dataset. (a) Representative histology image depicting HE staining of human DLPFC cross-section. (b) Scatterplot displaying hexagonally-packed locations of corresponding Visium spot centroids colored according to AAR annotation. (c) Three representative image patches from each AAR, each displaying a (185*μm*)^2^ region. (d) Adjacency matrix displaying the frequency with which patches of class [row] neighbor patches of class [column]. The number of instances of each class in the dataset is indicated in parentheses next to each row label in (d), as well as being displayed graphically in (e).

Annotation files for each tissue section specifying the spatial location of each foreground spot, along with manually-assigned layer annotation, were obtained from Maynard et al. [2020]. Using these foreground ST spot locations as centroids, patches were sampled to cover a 256 × 256 pixel region, which corresponded to a physical area 185 μm × 185 μm in each tissue. All 3-channel image patches were normalized such that *μ_RGB_* = (0.485, 0.456, 0.406), *σ_RGB_* = (0.229, 0.224, 0.225). The complete dataset consisted of 47,329 foreground patches with 7 unique classes across the 12 arrays. The number of patches belonging to each class is detailed in Fig. 4(d).

For experiments in this paper, the data were divided into six equally-sized folds for nested cross validation. The following fold compositions were chosen, using sample numbers from the original publication: Fold 1=(151507, 151508), Fold 2=(151509, 151510), Fold 3=(151669, 151670), Fold 4=(151671, 151672), Fold 5=(151673, 151674), Fold 6=(151675, 151676).

#### 2.1.4 Task Specification

As the data being considered in this paper are sampled according to regular grids — either cartesian or hexagonal — we represent inputs to the registration model as a tensors of dimension (*H_ST_*, *W_ST_*, *H_p_*, *W_p_*, *C*), where *H_ST_* and *W_ST_* represent the height and width of the ST grid, *H_p_* and *W_p_* represent the height and width of the patches, and *C* represents the number of image channels. As all data considered in this paper are square patches sampled from RGB images, we hold *H_p_* = *W_p_* and *C* = 3.

Labels are represented as 2D tensors of dimension (*H_ST_*, *W_ST_*). Thus, for input *X* and label *Y*, *X_i,j_* yields the image patch (of dimension *H_p_* × *H_p_* × 3) corresponding to row *i*, column *j* in the ST array, while *Y_i,j_* yields the class label for that patch. All foreground patches are assigned a label between 1 and *N*_class_ — where *N*_class_ represents the number of distinct foreground classes — while background patches are represented as zero-valued arrays and assigned a label of 0.

### 2.2 Model Specification

In this section, we discuss several variants of the proposed “GridNet” architecture for tissue registration, which is comprised of two component networks — a patch classification CNN and a segmentation CNN — that are connected and trained in an end-to-end manner. We provide an outline of the operation of the network on inputs constructed as outlined in the previous section, and additionally introduce a purely segmentation-based approach to tissue registration that will serve as a baseline for comparison of GridNet’s performance.

#### 2.2.1 GridNet architectures

The GridNet model is comprised of two components CNNs: patch classifier *f* and global corrector *g*. *f* accepts inputs of dimension (*H_p_*, *H_p_*, 3) and outputs a vector of predictions of length *N*_class_:

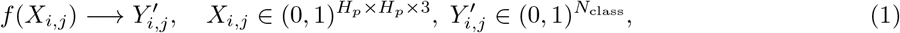

and *g* accepts inputs of dimension (*H_ST_*, *W_ST_*, *N*_class_) and outputs a matrix of the same dimension:

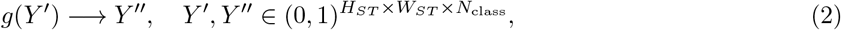

where *H_ST_* and *W_ST_* are the height and width (in patches) of the ST array, respectively.

The forward pass of the model *g* operates as follows: an input batch of shape (*B*, *H_ST_*, *W_ST_*, *H_p_*, *H_p_*, 3) (where B is the batch size) is flattened to a list of patches of dimension (*B* · *H_ST_* · *W_ST_*, *H_p_*, *H_p_*, 3). Each patch 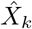 in this list is then passed through the patch classifier *f*, yielding an initial annotation 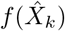. The resulting tensor of dimension (*B* · *H_ST_* · *W_ST_*, *N*_class_) is then reshaped to a tensor of dimension (*B*, *H_ST_*, *W_ST_*, *N*_class_). This tensor is then passed through the *g* network to obtain the final, identically-sized registration map. Prior to output and error calculation, all background patches {*X_i,j_* s.t. max(*X_i,j_*) = 0} are assigned class 0, while foreground patches are given a classification between 1 and *N*_class_. By design, GridNet cannot mistakenly classify background patches as foreground or vice-versa.

If the size of the input array is prohibitive for standard training by back-propagation, in which all intermediate activation states are stored in memory, the following strategy for gradient checkpointing is adopted: after obtaining the flattened patch list 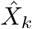, 0 < *k* ≤ (*B* · *H_ST_* · *W_ST_*),

1. Forward computation of mini-batch 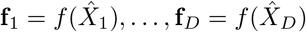. Save **f**_1_, …, **f***_D_*.
2. Repeat for all *D*-sized mini-batches 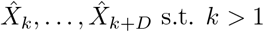, *k* + *D* ≤ (*B* · *H_ST_* · *W_ST_*).
3. Forward computation of *Y* = *g*(**f**_1_, …, **f***_B·H_ST_·W_ST__*).
4. Backward computation of the gradient to update parameters of *g*.
5. Forward computation of mini-batch 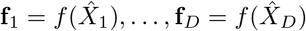.
6. Backward computation of the gradient to update parameters in *f* using only mini-batch 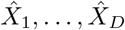. Save the gradients.
7. Repeat previous two steps for all *D*-sized mini-batches 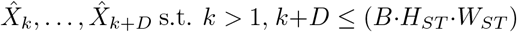.
8. Average gradients for all **f**_1_, …, **f***_B·H_ST_·W_ST__* and apply parameter update.

This methodology effectively partitions the input array into mini-batches of size *D*, where *D* can be chosen based on the size of the input arrays and available GPU memory. For all experiments in this paper using sufficiently large inputs (*H_p_* = 256), *D* = 32 was chosen as the mini-batch size after experimentation on NVidia V100 32GB GPUs.

Figure 5 details several variants of the GridNet architecture, defined by their choices for *f* and *g*. In GridNetSimple, a modified version of ResNet18 [He et al., 2016] is employed for *f*, in which batch normalization layers are removed and the number of filters in each convolutional layer is reduced by a factor of four. In the more sophisticated GridNet and GridNetHex models, DenseNet-121 [Huang et al., 2016] is employed for *f* instead. The choice of architecture for *g* is dependent upon the data being considered. When performing registration on either the ABA or Maniatis datasets, in which image patches are sampled according to a Cartesian grid, standard 2D convolutional kernels are employed to update the classification of a patch based on the classification of its neighbors. When considering the hexagonally-sampled Maynard dataset, specially-formulated hexagonal kernels are employed instead to accurately capture the 6-neighborhood around each spot. In this study, this was accomplished through the use of hexagonal convolution operations as implemented in the HexagDLy [Steppa and Holch, 2019] extension to PyTorch.

**Figure 5:**
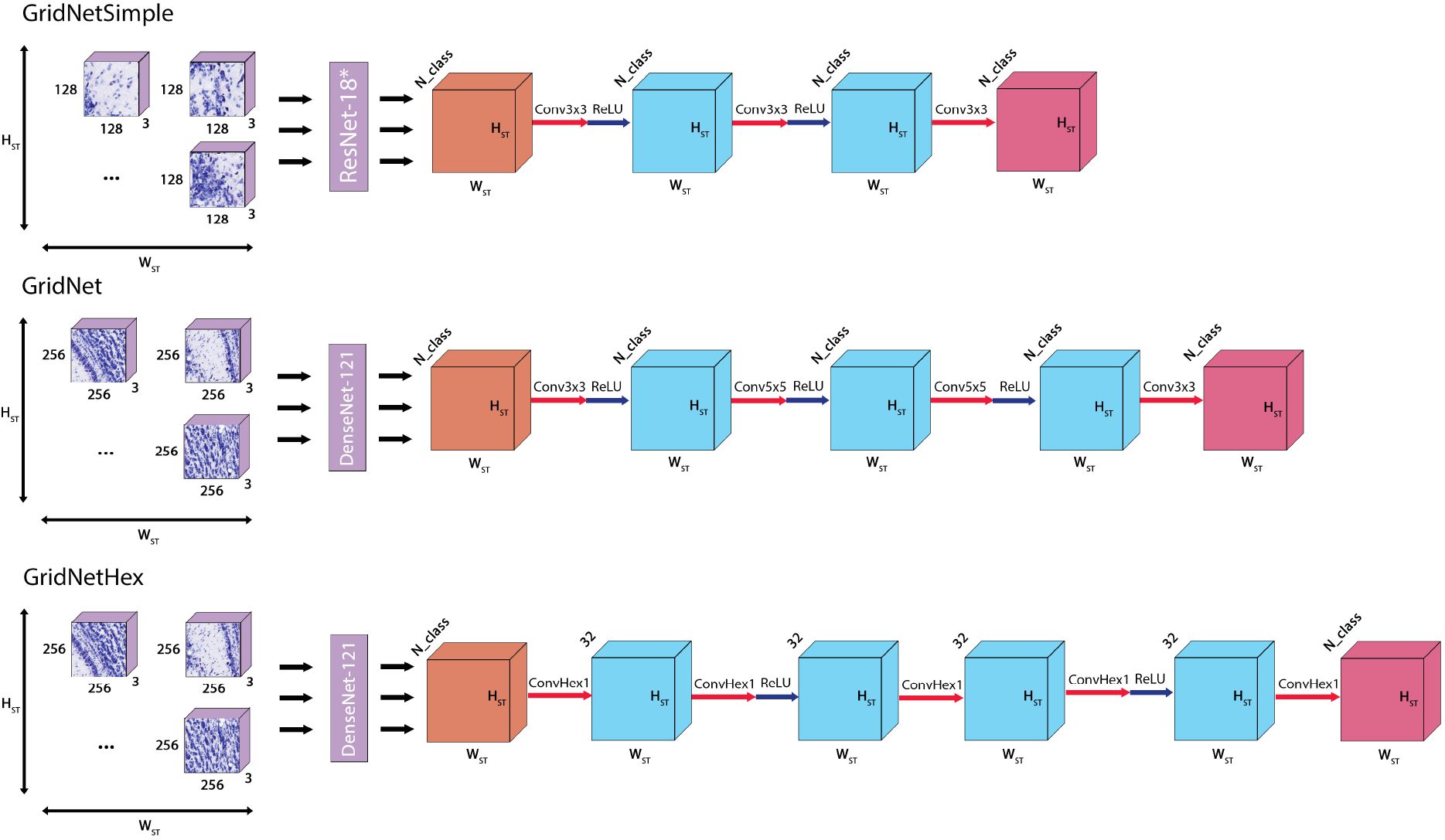
Three variant convolutional model architectures employed in image registration. ResNet18* indicates the modified ResNet architecture described in Section 2.2. Convkx*k* layers indicate 2-dimensional standard convolutional layers with kernel size *k*, stride-1, and same-padding. ConvHex*k* layers indicate 2-dimensional hexagonal convolutional layers with kernel size 1, stride-1, and same-padding. ReLU arrows indicate a rectified linear unit activation function.

#### 2.2.2 Segmentation architectures

As a baseline for comparison of the registration accuracy of GridNet, we formulated a modified version of ResNet-34 adapted to the registration task presented in this paper (Supplementary Fig. 1). Inputs to this network were transformed from the inputs to GridNet, flattened along each ST dimension to yield a single “stitched” image of dimension (*H_ST_* · *H_p_*, *W_ST_* · *H_p_*, 3).

The ResNet-34 architecture was modified in two significant ways in order to process these data. Firstly, the fully-connected layers were discarded, and the final 2D adaptive average pooling layer was modified to output tensors of dimension (*H_ST_*, *W_ST_*, *N_f_*), where *N_f_* is the number of filters at this layer in the network. A final 1×1 2D convolution with stride 1, same-padding, and *N*_class_ output filters was applied to this intermediate, yielding an output of dimension (*H_ST_*, *W_ST_*, *N*_class_). This enabled us to train the model on the same inputs and outputs as GridNet. The second modification was the reduction of the number of filters in each layer by a factor of four, which was required to fit even a single array from the ABA dataset into memory during training on a 32GB NVIDIA V100 GPU. As we had difficulty fitting even such a simple network in the memory of the GPU, we did not use U-Net style networks Ronneberger et al. [2015], which would require more memory.

### 2.3 Training Regimens

For all models discussed in this study, the optimization criterion is the cross-entropy loss between the predicted class probabilities and their true labels.

When training the *f* network alone, the problem reduces to simple image classification. For an input batch of image patches 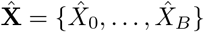 (where all 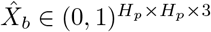) and associated labels 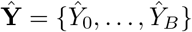 (where all 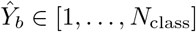), the objective function is given as:

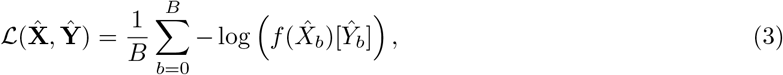

where 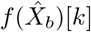 indicates the predicted probability of patch *b* belonging to class *k*, and 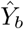 indicates the index of the true class of patch *b*.

When training the *g* network alone, or both networks together, image patches and their tensors are arranged into two-dimensional tensors as described in the previous section. For an input batch of image patch arrays **X** = {*X*_0_, …, *X_B_*} (where all *X_b_* ∈ (01),^*H_ST_*×*W_ST_*×*H_p_*×H_p_×3^) and their associated labels **Y** = {*Y*_0_, …, *Y_B_*} (where all *Y_b_* ∈ [1,…, *N*_class_]*^H_ST_×W_ST_^*), the objective function is given as:

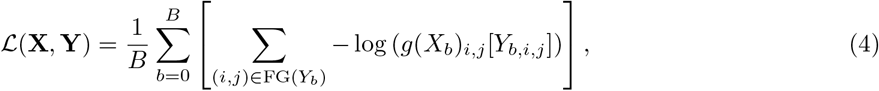

where FG(*Y_b_*) indicates the set of foreground patch coordinates in array *b*, *g*(*X_b_*)*_i,j_* [*k*] indicates the predicted probability of the patch at location (*i*, *j*) in array *b* belonging to class *k*, and *Y_b,i,j_* indicates the index of the true class of the patch at location (*i, j*) in array *b*.

Three competing training strategies were investigated for experiments on GridNet, all employing the Adam optimizer:

#### Two-stage training

1. Train *f* on the set of all foreground patches in the training set using batch size *B* · *H_ST_* · *W_ST_* patches for *E*_1_ epochs.
2. Fix parameters of *f*.
3. Train *g* using a batch size of *B* image arrays for *E*_2_ epochs.
4. Return the best-performing model on the validation set.

#### At-once training

1. Train *f* and *g* simultaneously, using learning rate of *lr* for *g* and *α* · *lr* for *f*, using a batch size of *B* image arrays for *E* epochs.
2. Return the best-performing model on the validation set.

#### Fine-tuning training

1. Initialize *f* and *g* with parameter values obtained from two-stage training approach. Save the state of the optimizer for the parameters in *g*.
2. Load the state of the optimizer for the parameters in *g*, and re-initialize the optimizer for *f*. Train *f* and *g* simultaneously, using learning rate of *lr* for *g* and *α* · *lr* for *f*, using a batch size of *B* image arrays for *E*_3_ epochs.
3. Return the best-performing model on the validation set.

### 2.4 Implementation Details

All code is written in PyTorch [Paszke et al., 2019] and is publicly available at https://github.com/flatironinstitute/st_gridnet/.

## 3 Results

In this section, we investigate several regimens for training the proposed GridNet model, then compare the registration performance attained by our model against competing approaches. We provide a schematic illustration of GridNet in Fig. 1b, as well as summary figures of the three datasets considered in Figs. 2, 3 and 4.

### 3.1 Determining optimal training strategy

Due to the complexity of the GridNet architecture, we began by exploring training procedures for the network. For this analysis, we considered a simplified form of the architecture, GridNetSimple (Section 2.2, Fig 5), which removes batch normalization layers from the patch classifier *f* (see Ioffe and Szegedy [2015] for a discussion of batch normalization in neural network training) in order to account for the different effective batch sizes of *f* and *g*. We trained the GridNetSimple architecture using the ABA reference histology dataset (Section 2.1.1, Fig. 2) with a patch size of 128 pixels. This smaller patch size was chosen in order to limit the size of input images and fit a batch size of *B* = 2 arrays into GPU memory during training.

#### 3.1.1 Two-stage training

We first considered a two-stage training approach, in which we first train *f* alone on the set of all image patches in the training set for *E*_1_ = 50 epochs, then fix the parameters of *f* and train *g* for *E*_2_ = 50 epochs. Initially, we employed the same learning rate for both training phases, and repeated the fitting for ten random samples from the interval log(*lr*) ~ Uniform(−4, −3) (Fig. 6a). During these experiments, we ensured that training batches for *f* and *g* contained the same number of total image patches in order to remove batch information content as a confounding factor. In Fig. 6a, we see that the best model obtained after training *f* alone attains 59% registration accuracy on the validation set, while the best model obtained after training *g* using fixed *f* attains 84% registration accuracy on the same validation set. We note that this large increase in validation set performance is observed whether the weights of *f* are initialized randomly or by pre-training on a corpus such as ImageNet [Deng et al., 2009]. Such pre-training merely reduces the number of training epochs required to attain the same performance in *f* (Supplementary Fig. 2). This finding is consistent with those in Raghu et al. [2019], who determined that transfer learning from ImageNet is more important for the correct initialization of the *magnitudes* of medical image network parameters, rather than their specific values. As a result, and due to the relatively large scale of the ABA dataset, we relied on models trained from randomly initialized weights for the remainder of this study.

**Figure 6:**
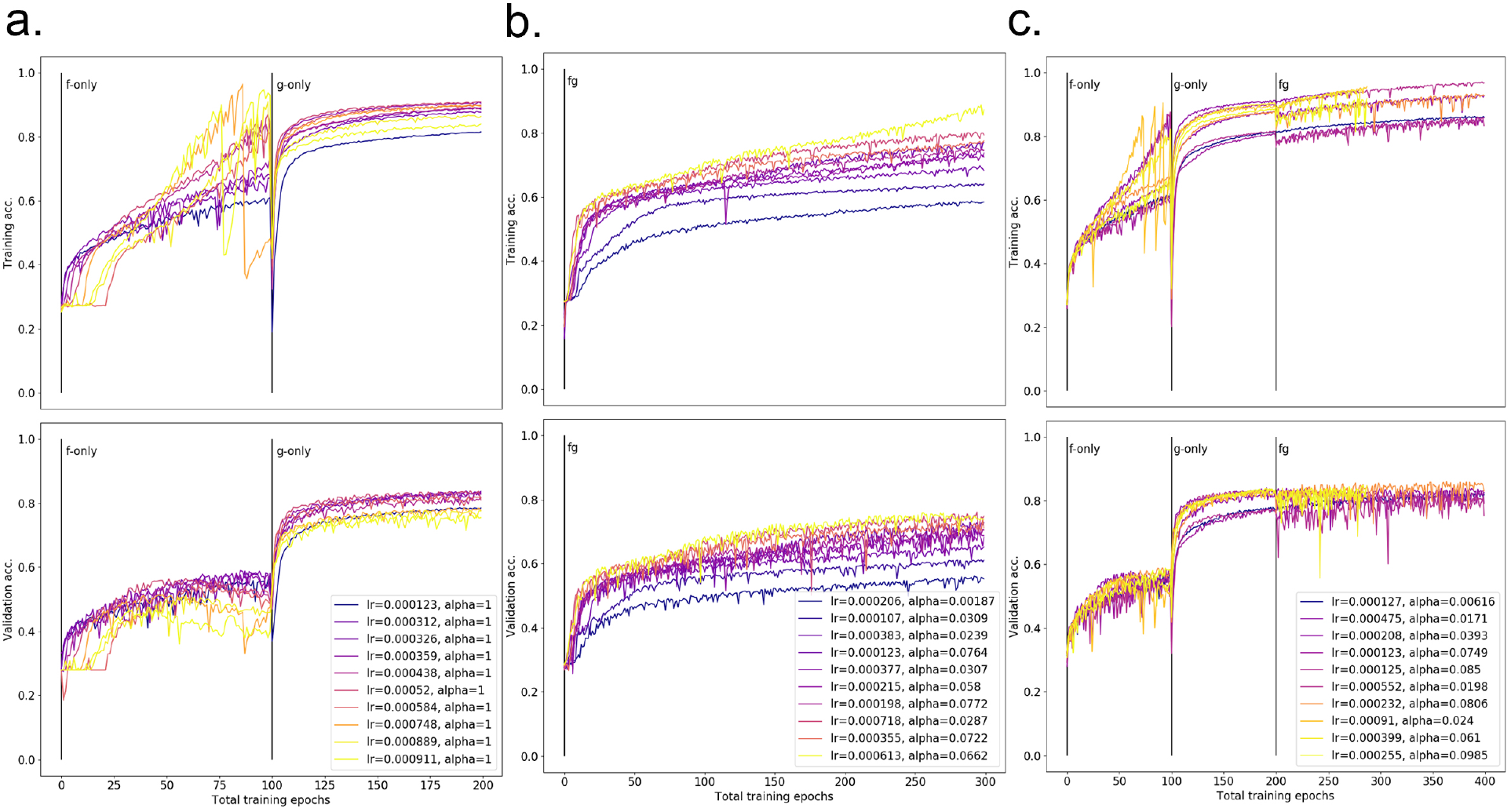
Registration accuracy attained by GridNetSimple architecture on ABA data under (a) two-stage (b) at-once and (c) fine-tuning training regimens (see Section 2.3). Each sub-plot shows training (top) and validation (bottom) accuracy as a function of training epoch, with a separate trace for each of 10 randomly sampled learning rates. Across all plots, “lr” denotes learning rate for parameters of *g*, and “alpha” denotes fraction of lr used as learning rate for parameters of *g*. Individual traces are color-coded from cool to warm according to the value of log(lr · *α*).

#### 3.1.2 End-to-end training

While the results from the two-stage approach are encouraging, demonstrating the power of this hybrid classification approach even when both component models are extremely simple, training both components simultaneously is conceptually more appealing. Such an “end-to-end” training approach would allow local learning rules to be updated in response to observed global patterns.

With this in mind, we next attempted training all model parameters at once from randomly initialized weights for *E* = 300 epochs. While the parameters from *f* and *g* would be updated simultaneously under this regimen, it merited further consideration as to whether their parameters should change at the same rate. In the two-stage regimen, we held the learning rate for *f* and *g* to be equal for simplicity, but observe that higher learning rates lead to strong fluctuations in both training and validation set performance during optimization *f* (Fig. 6a). We hypothesized that this was due to the fact that parameters of *f* are coupled across all input image patches, thus impacting model predictions disproportionately when stepped at the same rate as the parameters of *g*. To address this, we allowed for the learning rate of *f* to be set at some fraction *α* of the learning rate for *g* during training. We found that this strategy gave the best results when *α* < 0.1, (Supplementary Fig. 3), and as such sampled *α* ~ Uniform(0, 0.1) in subsequent analyses.

Accounting for this difference in learning rate, we see in Fig. 6(b) that the best performing model trained in the at-once regimen (76% validation accuracy) still fails to match the validation set accuracy attained by the two-stage model, despite surpassing the accuracy attained by training *f* alone. In order to better guide the joint optimization in this large parameter space, we considered a new “fine-tuning” approach to training, in which *f* and *g* are pre-trained following the two-stage regimen (*E*_1_ = *E*_2_ = 50 epochs), then jointly trained as in the at-once approach for an additional *E*_3_ = 100 epochs. In Fig. 6(c), we see that the best-performing model under this training regimen (86% validation accuracy) slightly outperforms the top models obtained by the two-stage approach, indicating the model may correct some errors arising from performing classification and segmentation in disjoint steps. While the at-once approach is appealing due to its simplicity, the relative success of the fine-tuning approach suggests that such an optimization is difficult even for simple models, and proper initialization of the parameters of *f* and *g* will be vital for success.

### 3.2 Registration of Cartesian ST data

Encouraged by our findings in the previous section, we increased the complexity of both *f* and *g* and applied the GridNet architecture (Section 2.2, Fig. 5) to both the ABA and the Maniatis datasets. For these analyses, we consider wider image patches (256 pixels) so that we may provide more information to our patch classification network and investigate the added benefit of the global segmentation layer under these conditions.

This increase in input size (≈ 1GB per tissue in each dataset) quickly exhausted the limits of memory on an NVidia V100 32GB RAM GPU during training with back-propagation, even when employing a batch size of just one array. In order to accommodate these large data, we implemented *gradient checkpointing*, described in detail Section 2.2.2., which allows the model to process input arrays in GPU-friendly minibatches, and enables end-to-end training in instances where input arrays prohibitively large for standard back-propagation. In order to simulate batch sizes *greater* than one, we additionally implemented a *gradient accumulation* step, in which back-propagated gradients are summed over a number of input batches (five, in our experiments) before updating model parameters. This allows us to reduce noise in the parameter update step and aids in convergence to a local optimum.

We then sought to compare the performance of the improved GridNet against approaches based purely on classification or segmentation. For a pure classification approach, we applied the *f* network from Grid-Net (DenseNet-121) to all foreground image patches in the dataset independently. For a pure segmentation approach, we employed “ResNet-34-Seg,” a ResNet-34 architecture modified to produce registration maps from whole-slide images (see Section 2.2.3). Due to the size of the images, the number of filters in each layer of ResNet-34-Seg had to be reduced by a factor of four in order to accommodate a single input during training on a 32GB RAM GPU. For each model and dataset combination, we performed hyperparameter optimization by sampling 10 learning rates (and *α*’s, if applicable) according to log(lr) ~ Uniform(−4, −3), *α* ~ Uniform(0, 0.1). From the five best-performing models on the validation set (Supplementary Table 1), we constructed an ensemble classifier by unweighted voting to yield an estimate of best-case performance. We additionally calculated the mean and standard deviation of accuracy and area under receiver operator curve (AUROC) across the component models to yield an estimate average-case performance.

In Tables 1 (ABA) and 2 (Maniatis), we see that across both datasets, GridNet greatly outperforms both DenseNet-121 and ResNet-34-Seg in average case and best case registration accuracy, per-class AUROC, and macro average AUROC. Between the competing approaches, we see that registration by ResNet-34-Seg performs worse than registration by DenseNet-121 along the same metrics for both datasets, despite the fact that DenseNet-121 only operates on dissociated patches. We attribute this to the reduction in complexity of the ResNet-34-Seg architecture necessitated by the size of input images, and highlights the difficulty of performing segmentation for images of this size. We additionally note that the patch classification accuracy attained by DenseNet-121 is much lower in the Maniatis dataset than the ABA dataset. This is likely due to properties of the data themselves: the ABA dataset is a well-curated reference histology dataset, with highly consistent staining and sample orientation. The Maniatis dataset, on the other hand, presents substantial variability in both tissue orientation and stain intensity, both of which can be confounding to image classification architectures. The predictive power of DenseNet may be enhanced by preprocessing images with stain normalization techniques, and additionally by the application of data augmentation to introduce robustness to orientation. Despite this variability, we see that the segmentation layer of GridNet adds substantially to the generalized registration accuracy on the Maniatis dataset (almost 12%), indicating that this approach is beneficial even when built on top of an imperfect classifier.

**Table 1:** Accuracy and AUROC attained by registration models on ABA independent test set. Results shown were calculated using the five best-performing models obtained during hyperparameter optimization using the validation set. “Ensemble” denotes results obtained by ensemble classifier built by unweighted voting of the five best models, while mean and standard deviation (SD) for performance metrics are calculated using the independent predictions from said models. AUROC is reported for each class (one-vs-rest) and for macro average across all 13 classes.

|  |  | Densenet-121 |  | GridNet |  | ResNet-34-Seg |  |
| --- | --- | --- | --- | --- | --- | --- | --- |
|  |  | Ensemble | Mean (SD) | Ensemble | Mean (SD) | Ensemble | Mean (SD) |
| Accuracy |  | 0.861 | 0.826 (4.4e-3) | 0.912 | 0.892 (3.7e-3) | 0.731 | 0.678 (3.08e-2) |
| AUROC | 1. Midbrain | 0.988 | 0.981 (2.30e-3) | 0.996 | 0.994 (8.79e-4) | 0.935 | 0.921 (1.91e-2) |
|  | 2. Isocortex | 0.988 | 0.985 (7.34e-4) | 0.997 | 0.995 (1.27e-3) | 0.980 | 0.970 (5.79e-3) |
|  | 3. Medulla | 0.994 | 0.992 (1.37e-3) | 0.998 | 0.998 (1.10e-3) | 0.959 | 0.942 (8.00e-3) |
|  | 4. Striatum | 0.978 | 0.971 (1.40e-3) | 0.991 | 0.989 (2.61e-3) | 0.943 | 0.915 (2.04e-2) |
|  | 5. Cerebellar nuclei | 0.998 | 0.996 (1.32e-3) | 0.999 | 0.998 (1.25e-3) | 0.980 | 0.959 (9.38e-3) |
|  | 6. Cerebellar cortex | 0.998 | 0.997 (2.71e-4) | 0.999 | 0.999 (2.08e-4) | 0.985 | 0.978 (2.93e-3) |
|  | 7. Thalamus | 0.994 | 0.990 (1.33e-3) | 0.998 | 0.997 (7.22e-4) | 0.966 | 0.948 (1.36e-2) |
|  | 8. Olfactory areas | 0.986 | 0.980 (7.38e-4) | 0.994 | 0.993 (1.20e-3) | 0.971 | 0.953 (3.93e-3) |
|  | 9. Cortical subplate | 0.971 | 0.962 (3.22e-3) | 0.983 | 0.981 (1.27e-3) | 0.935 | 0.894 (1.98e-2) |
|  | 10. Pons | 0.989 | 0.982 (4.08e03) | 0.996 | 0.995 (1.10e-3) | 0.930 | 0.908 (1.36e-2) |
|  | 11. Pallidum | 0.973 | 0.964 (4.71e-3) | 0.988 | 0.984 (2.62e-3) | 0.931 | 0.897 (1.36e-2) |
|  | 12. Hippocampal formation | 0.991 | 0.987 (6.55e-4) | 0.997 | 0.995 (1.25e-3) | 0.977 | 0.960 (8.81e-3) |
|  | 13. Hypothalamus | 0.992 | 0.987 (1.47e-3) | 0.997 | 0.995 (4.77e-4) | 0.947 | 0.928 (1.56e-2) |
|  | Macro Average | 0.988 | 0.983 (1.09e-2) | 0.995 | 0.993 (5.4e-3) | 0.957 | 0.937 (2.99e-2) |

**Table 2:**
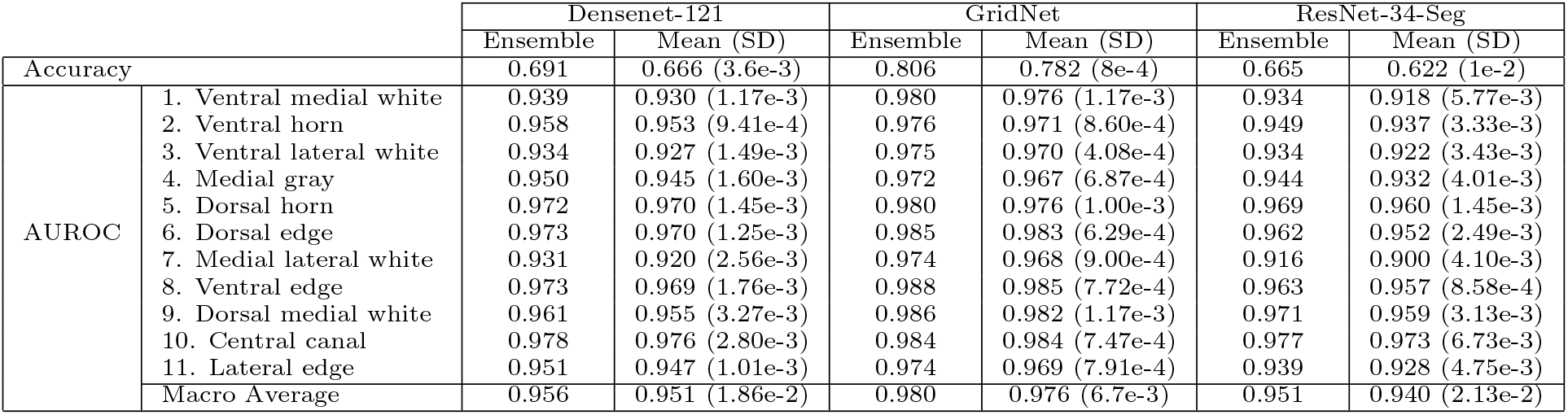
Accuracy and AUROC attained by registration models on Maniatis independent test set. Results shown were calculated using the five best-performing models obtained during hyperparameter optimization using the validation set. “Ensemble” denotes results obtained by ensemble classifier built by unweighted voting of the five best models, while mean and standard deviation (SD) for performance metrics are calculated using the independent predictions from said models. AUROC is reported for each class (one-vs-rest) and for macro average across all 11 classes.

We note that in both datasets, GridNet displays uniformly high AUROC for one-vs-rest (OVR) class predictors (Tables 1, 2), yet may fail to achieve a similarly impressive level of registration accuracy. To understand this, we generated confusion matrices for each dataset to get a more granular view on the types of errors being made by GridNet (Fig. 7). In both ABA and Maniatis datasets, we notice that there is substantial overlap between the large off-diagonal elements of the confusion matrices, which indicate persistent errors mistaking one class for another, and the large off-diagonal elements in the class adjacency matrices (Figs. 2d, 3d). This suggests that many of the errors made by GridNet involve mistakes between classes that are frequently neighbors, such as pallidum and striatum in ABA, and central canal and medial gray matter in Maniatis. Furthermore, we note that GridNet most frequently misclassifies patches belonging to rare classes, such as cerebellar nuclei, cortical subplate, and pallidum in ABA, and central canal and lateral edge in Maniatis. This suggests that in such boundary cases, GridNet may default to making predictions close to the class prior (the frequency of each class in the training data) in order to minimize the chance of making costly mistakes. This may partially explain the discrepancy between registration accuracy and AUROC observed in Tables 1 and 2, and suggests that AUROC may be a better reflection of the model’s performance.

**Figure 7:**
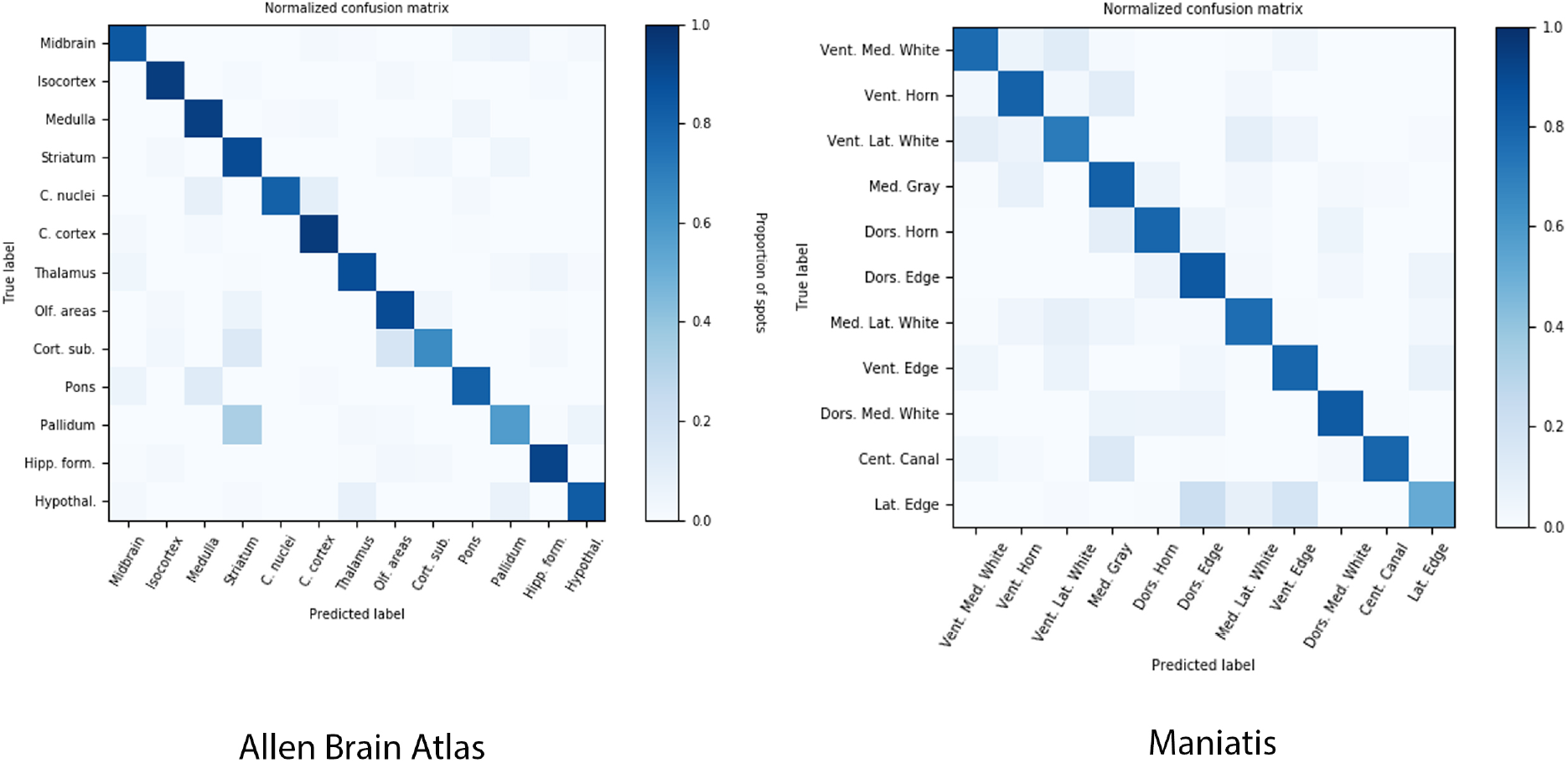
Confusion matrices for GridNet model on (left) ABA and (right) Maniatis datasets. Results shown were calculated on the held-out test set using the best-performing model on validation set. Rows are normalized such that each entry contains the proportion of patches with true label [row] that are classified as [column].

These prevalent boundary errors may be influenced by the presence of noisy labels, resulting from the inherent difficulty of drawing hard boundaries on continuously-varying tissue. In Fig. 8, we visualized the probability of misclassifying each patch — 1 – *P*(correct), where *P*(correct) is calculated using the classification probabilities output by the best-performing model and the index of the true label — across representative tissue arrays from each dataset. For predictions made using the *f*-network only, we see a rather uniform distribution of misclassification probability density, as well as a “spiking” behavior where certain patches are far more likely to be misclassified than their neighbors despite being located in regions of mono-class tissue. Using the full network, we see that the misclassification density is greatly reduced overall, and the remaining uncertainty is highly concentrated near boundaries between tissue classes, or boundaries between foreground and background patches. This suggests that GridNet is able to learn from the context of surrounding tissue to correct errors, and when properly trained, persistent error may simply reflect inescapable error in tissue labeling.

**Figure 8:**
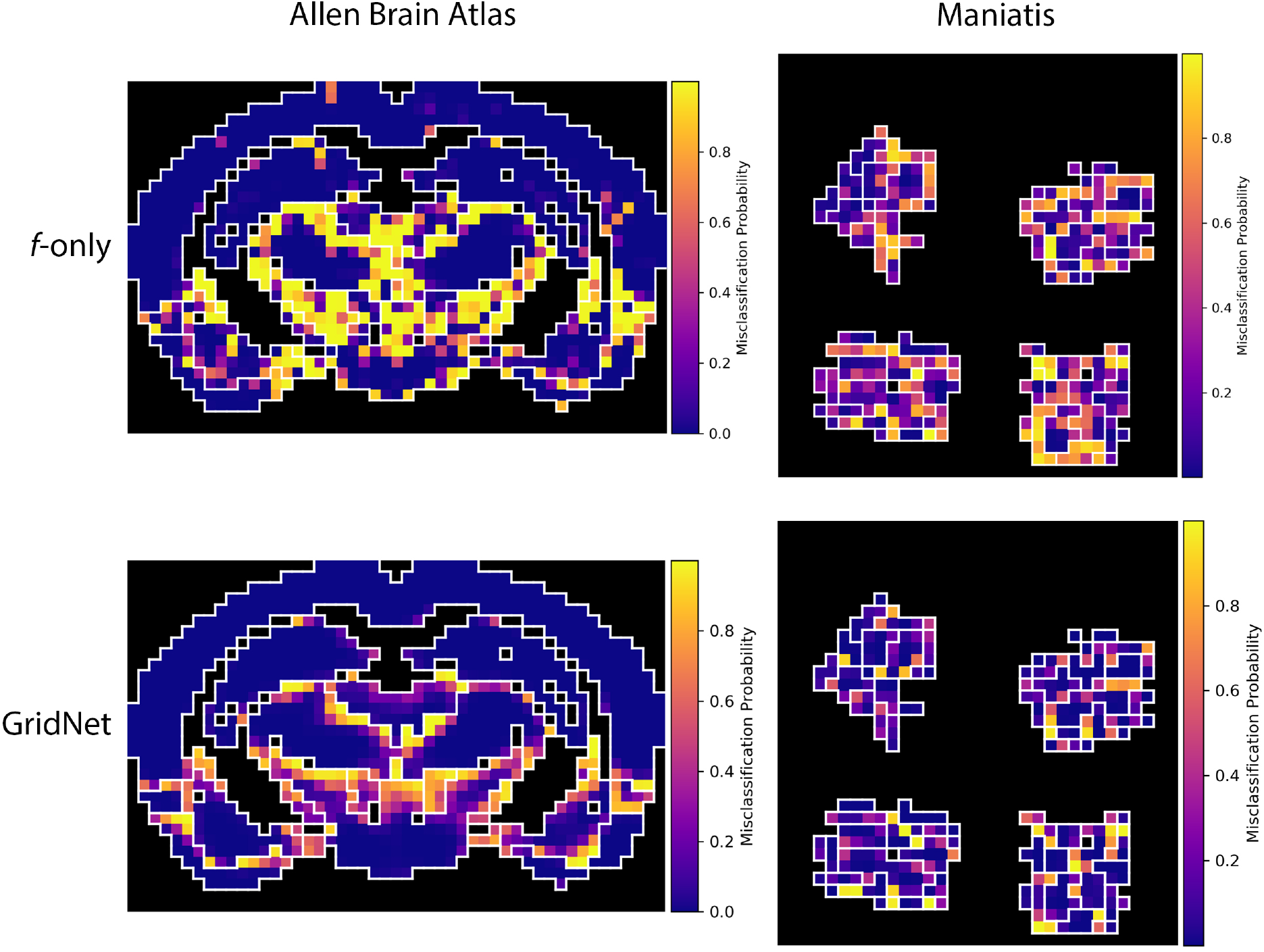
Misclassification probability density maps for representative images from (left) ABA and (right) Maniatis mouse spinal cord datasets. Each pixel is colored according to 1-P(correct), as estimated by either the full GridNet model (top row) or the *f* component only (bottom row), for the corresponding image patch. Patches indicating slide background are rendered in black, and class boundaries are denoted by white outlines.

### 3.3 Registration of non-Cartesian ST data

Finally, we sought to demonstrate the utility of GridNet for spatial data which are sampled in a non-cartesian grid — namely, the recently-released Visium ST platform from 10X Genomics. The hexagonal spot array used in this assay yields greater packing density (see Fig. 4b), increasing spatial resolution, but precludes the use of standard 2D-convolutional operations in *g*. Our GridNet approach can still be applied to these data through the modification of *g* to employ *hexagonal* convolution operations, which take into account the true 6-neighborhood of each spot (see Section 2.2.1). Using this modified form of GridNet, GridNetHex (Section 2.1, Fig 5), we assessed the registration performance on the Maynard human DLPFC Visium dataset (Fig. 4). While the increased resolution of the Visium arrays yields a corresponding increase in the number of patches (4,992 spots per array in Visium compared to 1,155 in standard ST), the small number of tissues processed by this study (12) limits the amount we can expect to learn from these data alone. As a result, we applied two-stage training with *E*_1_ = *E*_2_ = 100 training epochs in each phase to minimize over-fitting, and initialized the parameters of *f* with the optimal values obtained from our ABA experiment (Section 3.2) in order to speed convergence. Furthermore, for this analysis we chose to assess the generalized performance of our model using six-fold nested cross-validation, rather than employing a fixed train-validation-test partition as we have for larger datasets. During nested cross-validation analysis, each fold was held out as a test set exactly once, while hyperparameters were separately optimized for each fold by performing cross-validation on the remaining five folds. For each fold of the cross-validation analysis, five parameters were drawn from the ranges log(lr) ~ Uniform(−4, −3), *α* ~ Uniform(0, 0.1). The models with the best validation performance for each cross-validation fold were used to generate an ensemble predictor for the corresponding test fold.

In Table 3, we see that across the six folds, the ensemble network built from the full GridNetHex model yields small gains in registration accuracy over the ensemble network built from the *f*-network alone. The negligible magnitude of improvement over patch classification alone suggests, unsurprisingly, that more samples will be needed in order for the *g* network to learn global patterns that can refine local predictions. Additionally, we see that the full network ensemble achieves slightly lower macro average AUROC than the f-network ensemble, likely indicating over-fitting of the *g* network to the limited training data. The relatively low accuracy of the patch classifier can be understood by examining the quality of the Maynard data (Fig. 4), which were collected using a confocal microscope instead of the higher resolution slide scanner employed in the other datasets. This resulted in lower-resolution patch images in which few sub-cellular features besides the size and placement of nuclei can be visibly distinguished. With higher-resolution imaging, as well as expansion of the number of tissue being trained upon, we will be able to increase the generalized accuracy of our DLPFC registration network. Despite this, our preliminary analysis demonstrates how GridNet may be applied to the registration of data collected from the Visium platform.

**Table 3:**
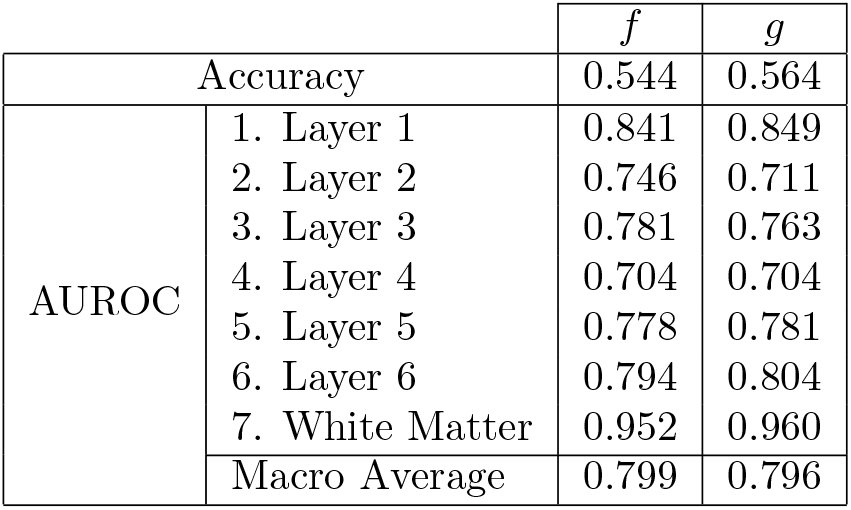
Performance of ensemble GridNetHex models on held-out test folds during nested cross-validation analysis of the Maynard dataset. Each column details the performance of an ensemble model created using the best-performing models found during cross-validation on the remaining five folds. Sub-columns distinguish between the performance of the ensemble created with the *f*-network only or the full model (*g*). AUROC is reported for each class (one-vs-rest) and for macro average across all seven classes.

## 4 Discussion

We present GridNet, a novel deep learning network for common-coordinate registration of high-resolution histology images. GridNet balances the need for sub-cellular resolution and awareness of spatial context by applying a hybrid classification-segmentation architecture to high-resolution image patches sampled sparsely across the tissue. Using this architecture, our models can predict tissue type for ROIs in large tissues while respecting the memory limits of commercially available GPUs.

Through experimentation on both simulated and experimental spatial datasets, we demonstrated that the hybrid classification-segmentation architecture consistently out-performed competing approaches in the registration of spatial transcriptomic data. Compared to a pure-segmentation approach based on ResNet-34 (which required substantial reduction in complexity in order to process the histology images being considered) and a pure-classification approach based on DenseNet-121, we saw significant reduction in error across the tissue, and notably a reduction in the chance of misclassifying tissue within a contiguous region. Most remaining error was concentrated on class boundaries, reflecting the difficulty of assigning discrete labels to a continuously-varying input. Near such boundaries, predictions made by GridNet or competing registration architectures may skew towards the class prior in an attempt to minimize average-case misclassification error. While this behavior may be expected even in the presence of perfect training labels, the presence of noisy labels may greatly exacerbate it. The incorporation of replicate labeling data gathered from independent pathologists may help in the reduction of labeling noise, though a robust solution to this problem will require modification of the training objective function to explicitly model label noise, rather than assuming a gold standard. Methods for training neural networks in the presence of noisy labels is an active area of research [Vahdat, 2017, Zhang and Sabuncu, 2018, Song et al., 2020], and may benefit future iterations of the GridNet model.

While the hybrid classification-segmentation approach is capable of attaining high registration accuracy, the complex nature of the network architecture demands specific consideration during training. We found that the coupling of parameters in the *f* network across all input image patches yielded a different ideal learning rate from *g*, adding an additional hyperparameter during model training that must be tuned to the data at hand. Furthermore, we determined that pre-training of both component networks prior to end-to-end fine-tuning is instrumental in attaining high performance. Through experimentation on the ABA dataset, we developed a two-stage pre-training approach that yielded consistently strong results: parameters of local predictor *f* are initialized with values that yield the highest performance on the stand-alone task of foreground patch classification, then parameters of the global corrector *g* are optimized using *f* as a fixed feature extractor. While this method was sufficient to produce models exceeding the performance of competing approaches, we hope that further study of the network properties will reveal end-to-end training strategies that are more robust to initial parameter values.

In our brief analysis of ST data gathered with 10x Genomics’ Visium platform, we demonstrated that the GridNet paradigm could be applied when patches are sampled in non-Cartesian grids. The logical extension is the application of GridNet to data in which patches are sampled in an irregular manner. In FISH-based ST data, for example, the regions of interest are not defined by the fixed grid capture probes, but instead by the locations of cell nuclei. One may imagine *f* being applied to classify extracted nuclei into discrete cell types, and *g* leveraging information from neighboring cells to correct said predictions. This could be accomplished by modeling nuclei as an undirected graph with edges weighted by Euclidean distance between them, and converting *g* to a graph convolutional neural network (GCNN) to aggregate information from neighbors. Such a formulation can be thought of as a super-set of GridNet, although standard or hexagonal convolutional operations should be used for *g* when possible due to their relative efficiency.

Finally, we are interested in developing future iterations of GridNet that leverage data from multiple modalities at once. As the ST experimental workflow produces paired measurements of both histology and gene expression, we may seek to combine information from these two modalities in our patch prediction network. A recent study by Tan et al. [2020] explored how integration of these two modalities within a single classifier may yield increased ability to predict tissue when considering ST spots independently, but did not consider how surrounding context could be used to refine these predictions. We believe that future iterations of GridNet could build off of such multi-modal feature extractors to build even stronger models for tissue registration. Furthermore, such multi-modal models could be adapted to the task of predicting one data modality from the other. Using a set of ST data, models could be trained to predict the expression of key genes from histology data alone, allowing researchers to predict expensive gene expression measurements from inexpensive histology images. Our architecture is well adapted for multi-modal contexts where orthogonal sensors, assays or tests are applied sparsely to a instance with a corresponding high resolution image. Here we focus on spatial genomics, but applications for such a context and resolution hierarchy abound. Thus, We believe the GridNet paradigm will be a useful and generalizable approach to making predictions on a variety of spatial biological data, and building upon the implementation presented in this paper, can scale with the increasing size and resolution of spatial data being acquired today.

## Footnotes

1 https://github.com/tare/span

2 http://help.brain-map.org/display/api/Image-to-Image+Synchronization

