## Supplementary Tables/Figures for "A convolutional neural network for common coordinate registration of high-resolution histology images"

### Supplementary Material: A convolutional neural network for common coordinate registration of high-resolution histopathology images

Aidan C. Daly*[1], Krzysztof J. Geras†[3,2], and Richard A. Bonneau‡[1]

[1]Center for Computational Biology, Flatiron Institute, U.S.A.
[2]Center for Data Science, New York University, U.S.A.
[3]NYU Grossman School of Medicine, U.S.A.

September 14, 2020

*
†
‡

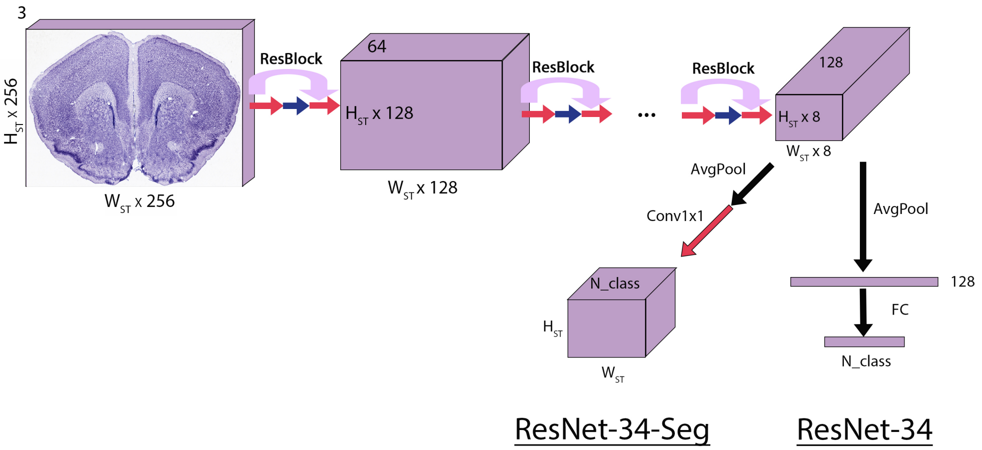


Figure 1: Modifications applied to ResNet-34 in order to create ResNet-34-Seg for common coordinate registration of high-resolution histopathology images. "ResBlock" connections indicate the residual block comprising ResNet, which contain skip connections in order to feed foward low-level image information. "AvgPool" arrows indicates average pooling layers, while "FC" arrows indicate fully-connected linear layers.

| Model | ABA | | Maniatis | |
|---|---|---|---|---|
| | lr | $\alpha$ | lr | $\alpha$ |
| GridNet | 1.857e-4 | 5.618e-2 | 6.685e-4 | 3.527e-2 |
| | 8.106e-4 | 5.251e-2 | 8.052e-4 | 6.921e-2 |
| | 3.054e-4 | 8.632e-2 | 4.127e-4 | 1.254e-2 |
| | 2.379e-4 | 6.889e-2 | 6.77e-4 | 9.795e-2 |
| | 2.389e-4 | 2.835e-2 | 3.33e-4 | 1.657e-2 |
| ResNet-34-Seg | 1.029e-4 | | 3.438e-4 | |
| | 2.559e-4 | | 1.256e-4 | |
| | 1.664e-4 | | 1.017e-4 | |
| | 2.579e-4 | | 4.724e-4 | |
| | 4.199e-4 | | 1.257e-4 | |

Table 1: Highest performing hyperparameter combinations from validation study of GridNet and ResNet-34-Seg on Allen Brain Atlas and Maniatis datasets.

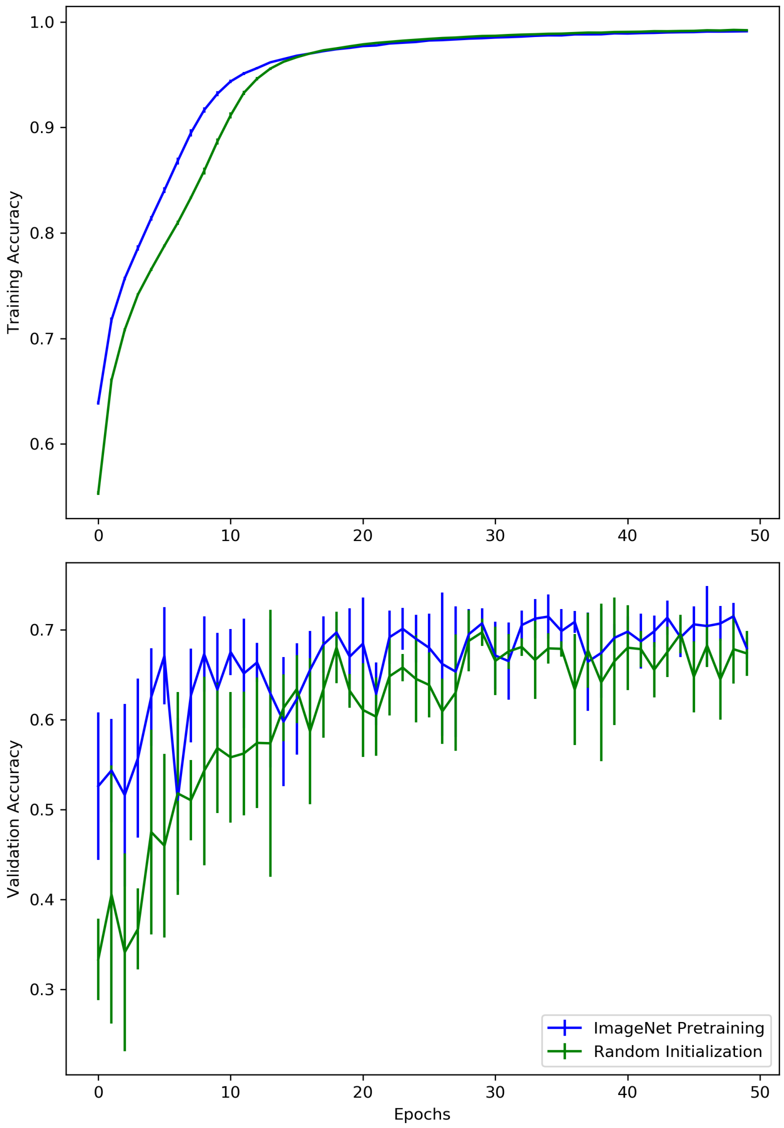


Figure 2: Registration accuracy attained by training ResNet-18 on image patches from the Allen Brain Atlas dataset under two training regimens: random weight initialization, and weight initialization by pre-training on ImageNet. Each training regimen was repeated five times using the Adam optimizer with a learning rate of 0.001. Mean training/validation accuracy is plotted for each regimen, with error bars denoting standard deviation.

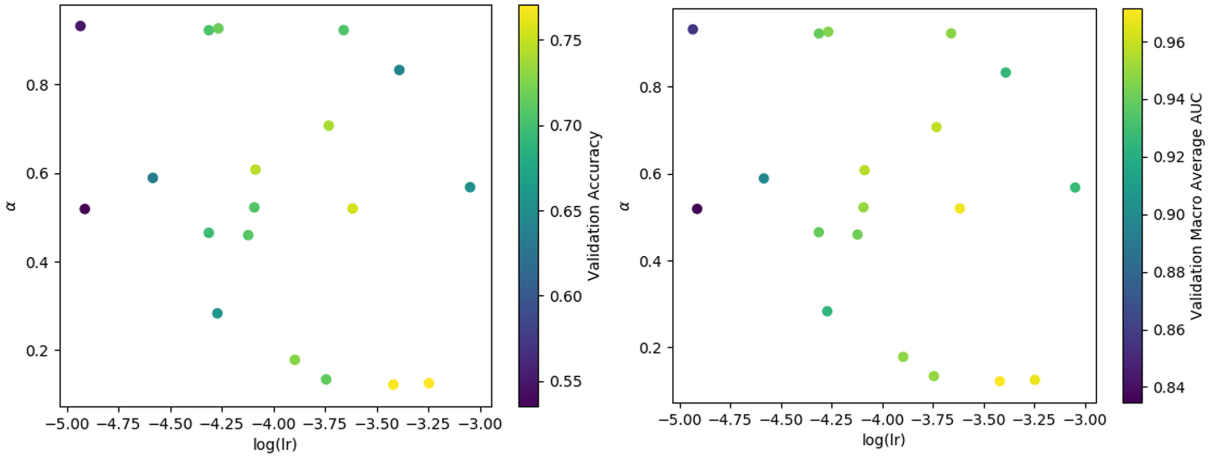


Figure 3: Effects of learning rate and alpha parameters on training of GridNetSimple registration model on Allen Brain Atlas dataset using the at-once training regimen. Each point represents a separate training from randomly initialized parameters, with points colored according to either the accuracy (left) or macro average ROC-AUC (right) attained on the validation set.
